# Influence of human activity on gut microbiota and immune responses of Darwin’s finches in the Galápagos Islands

**DOI:** 10.1101/2022.02.07.479384

**Authors:** Jada N. Bygrave, Maxine Zylberberg, Alyssa Addesso, Sarah A. Knutie

## Abstract

Urbanization can influence animal traits, including immunity and gut microbiota. Over the past several decades, the Galápagos Islands have seen rapid resident human population growth and tourist activity, leading to varying levels of human activity across Islands. Consequently, diet, gut microbiota, and immunity of endemic animals, such as Darwin’s finches, may have changed. The goal of this study was to determine the effect of land use on the immune response, gut microbiota, and body measurements of Darwin’s finches in 2008, at a time of rapidly increasing human activity in the Islands. Specifically, we compared proxies of immunity (lysozyme activity, and haptoglobin, complement antibody, and natural antibody levels), gut microbiota (bacterial diversity, community structure and membership, and relative abundance of bacterial taxa), and body measurements (body mass, tarsus length, and scaled mass index) across undeveloped, agricultural, and urban areas for medium ground finches (*Geospiza fortis*) and small ground finches (*G. fuliginosa*). We found that lysozyme activity was lower and observed bacterial species richness was higher in urban areas compared to non-urban areas across both finch species. For small ground finches, relative abundances of three bacterial genera (*Pseudoxanthomonas, Cloacibacterium*, and *Dietzia spp*.) were higher in urban areas compared to non-urban areas, but this pattern was not observed in medium ground finches. Medium ground finches were smaller in undeveloped areas compared to the other two areas, but body measurements of small ground finches did not differ across areas. Our results suggest that human activity can impact immune measures and gut microbiota of Darwin’s finches.

## INTRODUCTION

Human activity is rapidly changing the natural environment, which can influence avian ecology and evolution. For example, anthropogenic change can influence many avian traits, including morphology (e.g., bill size), behavior (e.g., boldness, predator avoidance, problem solving), and physiology (e.g., digestion, immunity) (Hendry *et al*., 2006; Møller, 2008; Atwell *et al*., 2012; Samia *et al*., 2017; Sol *et al*., 2018; Strandin *et al*., 2018). One factor that can mediate the effect of human activity on avian traits is changes in food availability. Humans can change food resource availability for birds by increasing the abundance of unnatural food through wild bird feeders, the disposal of human food waste into the environment, and by reducing the abundance of natural food items through environment changes (Murray *et al*., 2016; Bosse *et al*., 2017; Start *et al*., 2018). In turn, birds lacking natural food resources may be forced to invest less in maintaining body mass or physiological processes, such as immune responses, which can be energetically costly (Sheldon and Verhulst, 1996; Svensson *et al*., 1998; Lochmiller and Deerenberg, 2000; Demas, 2004; Sternberg *et al*., 2012; Cornet *et al*., 2014; Howick and Lazzaro, 2014).

In addition, human activity can alter avian traits through impacts on the gut microbiota, the community of symbiotic microbial species that live within the gut of their host (Grond *et al*., 2018). Different environments, including human-altered ones, contain different microbial communities that can colonize the gut of the host. As a result, the gut microbiota of the host can change in response to changes in land use, including urbanization (Phillips *et al*., 2018; Teyssier *et al*., 2018; Knutie *et al*., 2019; Berlow *et al*., 2021), and pollution of water and soil (Jin *et al*., 2017). In turn, the gut microbiota can interact with host cells to change host physiological traits, such as the immune system (reviewed in Round and Mazmanian, 2009); increased bacterial diversity and the presence of specific bacterial taxa can be related to increased resistance to virulent parasites and pathogens. As human activity increases, along with the concern over parasite-causing diseases in birds, disentangling these relationships is important to understand how novel stressors can influence birds and help aid in their management and conservation.

The Galápagos Islands of Ecuador are relatively pristine but are facing increasing human activity, which allows for the study of host-associated microbiota and immune systems in response to urbanization. Over the past 20 years, ecotourism and the permanent resident human population has grown rapidly on the Islands (Walsh and Mena 2016). For example, from 2007 to 2017, the number of permanent human residents increased from approximately 20,000 to 30,000 people and tourist numbers increased from 140,000 to 225,000 people (Watkins and Cruz, 2007; Walsh and Mena, 2016). Consequently, in areas with a human presence (e.g., towns and popular tourist sites), the natural habitat and diet of endemic species have been altered with the introduction of agricultural plants and human processed food (de León *et al*., 2018). Studies have shown that changes in habitat type can affect gut microbiota and metrics of immunity of Darwin’s finches (Zylberberg *et al*., 2013; Michel *et al*., 2018; Knutie *et al*., 2019; Loo *et al*., 2019). However, all studies on the gut microbiota of Darwin’s finches have occurred since 2016 and so we know little about how earlier years of human activity has influenced microbiota in Darwin’s finches. Furthermore, the effect of human-influenced land use on the interaction between gut microbiota and immunity has received little attention. A better understanding of human impacts on avian immune systems could be especially important because introduced parasites, such as avian pox and *Philornis downsi*, have been wreaking havoc on the survival of birds in the islands (Wikelski *et al*., 2004; Fessl *et al*., 2010; Koop *et al*., 2016).

The goal of this study was to determine the effect of land use on the immune response, gut microbiota, and body measurements of Darwin’s finches in 2008. Specifically, we characterized the gut microbiota (bacterial diversity, community structure and membership, and relative abundance of bacterial taxa), immune response (lysozyme activity, and haptoglobin, complement antibody, and natural antibody levels), and body measurements (body mass, tarsus length, and scaled mass index) in small ground finches (*Geospiza fuliginosa*) and medium ground finches (*G. fortis*) across three different habitat types in 2008 on Santa Cruz Island in the Galápagos. Specifically, we caught finches in three different areas that varied in human activity and presence including an area with: 1) little human activity and no permanent human population (hereon, undeveloped), 2) developed land for agricultural activities but only a small permanent human population (hereon, agricultural), and 3) developed land for a substantial, permanent human population (hereon, urban).

Previous studies have shown that the immune response of Darwin’s finches can vary temporally across land use types. For example, Zylberberg *et al*. (2013) found that finches living in agricultural and undeveloped areas had different immune responses across years, whereas finches in urban areas showed no change across years. One explanation is that food availability changed across years but not in all habitats. Variation in food availability could either directly affect the immune response or the gut microbiota, which in turn would affect the immune response. However, Zylberberg et al. (2013) did not examine the effect of land use on the immune response within each year. Thus, for the present study, we examined the impact of land use on four metrics of immunity: lysozyme activity, and complement antibody, natural antibody, and haptoglobin (using PIT54 acute phase proteins) levels. We chose these four aspects of the innate immune system because each plays an integral and complementary role in the first line of defense against novel pathogens (as previously discussed in Zylberberg *et al*., 2013). In short, the complement system of proteins activates the lysis of foreign cells, enhances antibody activity, and directly destroys viruses (Hirsch, 1982; Murphy *et al*., 2017). Natural antibodies bind novel pathogens, facilitate phagocytosis, and promote cell lysis (Casali and Schettino, 1996; Caroll and Prodeus, 1998). The PIT54 protein minimizes self-damage during inflammation and stimulates the white blood cell response upon pathogen exposure (Wicher and Fries, 2006; Quaye, 2008). Lysozymes lyse gram positive bacteria (Millet *et al*., 2007). Together, these measures afford a broad view of the innate immune system by providing information on both inducible and constitutive components of innate immunity.

We hypothesized that variation in land use would influence the immune response and gut microbiota of finches. Specifically, we predicted that finches in areas with lower human activity (agricultural and undeveloped areas) will have more similar immune responses and gut microbiota than finches living in areas with higher human activity (urban areas). Furthermore, because studies have shown that the metrics of the immune system can relate to changes in the gut microbiota community (Round and Mazmanian, 2009), we hypothesize that bacterial diversity will relate to immune measures in the finches. Finally, because food availability and thus diet differs across sites, we predict that body size (mass and tarsus length) and condition (scaled mass index; Peig and Green, 2009), will differ across land use type, as found in Knutie *et al*. (2019). Overall, our study will provide insight into the effect of human activity on the gut microbiota and immune proxies of island birds, which could have conservation implications for Darwin’s finches (Ohmer et al., 2021).

## METHODS

### Field site

We conducted this study on Santa Cruz Island, Galápagos, Ecuador during the breeding season in February 2008. To examine the effect of land use patterns on microbiota, immune response, and body size of finches, we sampled small ground finches and medium ground finches at six sites, spanning three different land use types (urban, agricultural, and undeveloped). We selected the study sites to represent the range of vegetation types and precipitation present in each land use and elevation category. The urban site was in Puerto Ayora (population < 20,000 in 2007; (Watkins and Cruz, 2007), in the lowland arid zone. Three sites were located within the agricultural zone of the island, which contains a combination of small farms, fruit plantations and cattle ranches. One agricultural site was in the lowlands and two in the highlands. Two sites were in undeveloped areas, representing both arid lowland and moist highland zones. The Galápagos is well known for extreme fluctuations in precipitation as a result of the El Niño weather pattern; however, 2008 was neither particularly wet nor particularly dry on average (Zylberberg *et al*., 2013).

### Sample collection

To obtain immune function and microbiota data, we caught birds in mist nets that were actively watched. Birds were removed from the mist net and processed as quickly as possible; in the event that multiple birds entered the mist net, they were removed and kept in cloth bags until they were banded and a blood sample collected. Body mass (g) and tarsus length (mm) were measured, and a scaled mass index (a metric of body condition) was calculated based on the methods from Peig and Green (2009). We banded each bird and collected blood samples using heparinized microcapillary tubes. For birds that defecated during handling, the fecal sample was immediately collected and placed in formalin until used for the fecal bacterial DNA extraction at the University of Connecticut in 2019. Blood samples were kept on ice (for 4-6 hours) until they were centrifuged to separate the red blood cells from the plasma. Plasma was then frozen at - 20°C.

### Immune measures

We used a hemolysis-hemagglutination assay to measure levels of natural antibodies (lysis activity) and complement antibodies (agglutination activity) (Matson *et al*., 2005). We used a commercial kit (from Tri-delta Diagnostics Inc., Morris plains, NJ, USA) to determine the plasma concentration of PIT54 acute phase protein (haptoglobin levels) following Millet *et al*. (2007). We used the lyso-plate assay described in Millet *et al*. (2007) to measure levels of lysozyme activity in plasma samples. Each of these assays was carried out with previously described protocols (Zylberberg *et al*., 2014).

### Bacterial DNA extraction and sequencing

Before starting the extraction from the finch feces, samples were centrifuged for 10 minutes at 10,000 rpm at 4°C and the supernatant (i.e., the formaldehyde) was then removed. Nanopure water (500 uL) was added, and the sample was vortexed and centrifuged again for 5 minutes at 4°C at 10,000 rpm. The supernatant was removed and the step was repeated. Nanopure water (200 uL) was then added to the sample and the sample was vortexed. Total DNA was extracted from finch feces using a Qiagen PowerFecal DNA Isolation Kit. The protocol listed in the kit was followed with the exception of the heating step; samples were heated for 30 minutes at 65°C. DNA extracts were then sent to the University of Connecticut Microbial Analysis, Resources and Services for sequencing with an Illumina MiSeq platform and v2 2×250 base pair kit (Illumina, Inc.). We also amplified a laboratory blank to control for kit contamination and had no detectable product. Bacterial inventories were conducted by amplifying the V4 region of the 16S rRNA gene using primers 515F and 806R (Caporaso *et al*., 2012) and with Illumina adapters and dual indices (Kozich *et al*., 2013). Raw sequences were demultiplexed with onboard bcl2fastq and then processed in Mothur v.1.42.3 (Schloss *et al*., 2009) according to the standard MiSeq protocol (Kozich *et al*., 2013). Briefly, forward and reverse sequences were merged. All sequences with any ambiguities, that did not align to the correct region, or that did not meet length expectations, were removed. Sequences were aligned to the Silva nr_v128 alignment (Quast *et al*., 2013). Chimeric reads were also removed using UCHIME (Edgar *et al*., 2011). Non-bacterial sequences that are classified as chloroplasts, mitochondria, or unknown (i.e., did not classify to the level of kingdom) were removed. Sequences were grouped into operational taxonomic units (OTUs) based on a 97% similarity level and identification of the OTUs was done using the Ribosomal Database Project Bayesian classifier (Wang *et al*., 2007) against the Silva nr_v128 taxonomy database. Alpha and beta diversity statistics were calculated by averaging 1,000 random subsampling of 4,000 reads per sample. We calculated observed species richness (sobs) and evenness, Shannon diversity index, and Simpson diversity index. Sobs describes the number of observed species and evenness describes the distribution of abundance across the species. The Shannon diversity index and Simpson index are estimators of species richness and species evenness but species richness is weighted more for Shannon and species evenness is weighted more for Simpson. The resulting data sets included a total of 1,827,365 sequences and an average of 25,032 reads per sample (min: 4,079, max: 121,506).

### Statistical analyses

Generalized linear models (GLMs) were used to determine the effect of land use on bacterial diversity metrics (sobs, Shannon Index, Simpson Index, and evenness), immune metrics (lysozyme activity, and haptoglobin, complement antibody, and natural antibody levels), and body size and condition (body mass, tarsus length, and scaled mass index). Shapiro-Wilks tests were used to determine whether the variables met normality; if the variables did not meet normality, they were log transformed for the analyses. Analyses were conducted using the glm function within the lme4 package (Bates *et al*., 2015). Probability values were calculated using log-likelihood ratio tests using the Anova function in the car package (Fox and Weisberg, 2019). Analyses were conducted in R (2021, version 4.0.4) and RStudio (2021, version 1.4.1103).

The effects of location and species on bacterial community dynamics were examined using the Bray-Curtis dissimilarity and Jaccard similarity matrices. The matrices were created using the ‘vegdist’ function in the vegan package in R (Oksanen *et al*., 2019). The ‘adonis2’ function in the vegan package was used to perform PERMANOVA to assess the differences in bacterial community composition and membership between different species and locations (Oksanen *et al*., 2019). Both Bray-Curtis and Jaccard are dissimilarity matrices, with Bray-Curtis taking into account the relative abundances of shared taxa (community structure) while Jaccard only considers the presence or absence of such taxa (community membership). Principal Coordinates Analyses (PCoA), where the distances among the samples are converted onto a graph, were done to compare and visualize differences between groups.

Relative abundances (arcsine square root transformed; (Shchipkova *et al*., 2010; Kumar *et al*., 2012) of bacterial phyla and genera were compared between species and location groups. Data were manipulated using packages tidyr, reshape2, and plyr in R (Wickham, 2007, 2011; Wickham and Henry, 2019) and ANOVAs were run in the car package in R (Fox and Weisberg, 2019); false discovery rate (FDR) tests were used to control for multiple analyses. A Tukey post-hoc test was done to determine which land use types differed from each other. All figures were created in Prism (2020, version 9).

## RESULTS

### Effect of land use and finch species on the gut microbiota

Bacterial diversity, as measured by sobs, differed among land use types for finches (Table S1; Fig. 1A). Birds in the urban area had higher sobs values than birds in agricultural and undeveloped areas (Table S2; *P* < 0.05). However, there was no effect of species or an interaction between species and land use type on sobs values (Table S1). Land use, bird species, and their interaction did not affect Shannon Index, Shannon evenness, and the Simpson Index (Tables S1-S2).

**Fig 1.**
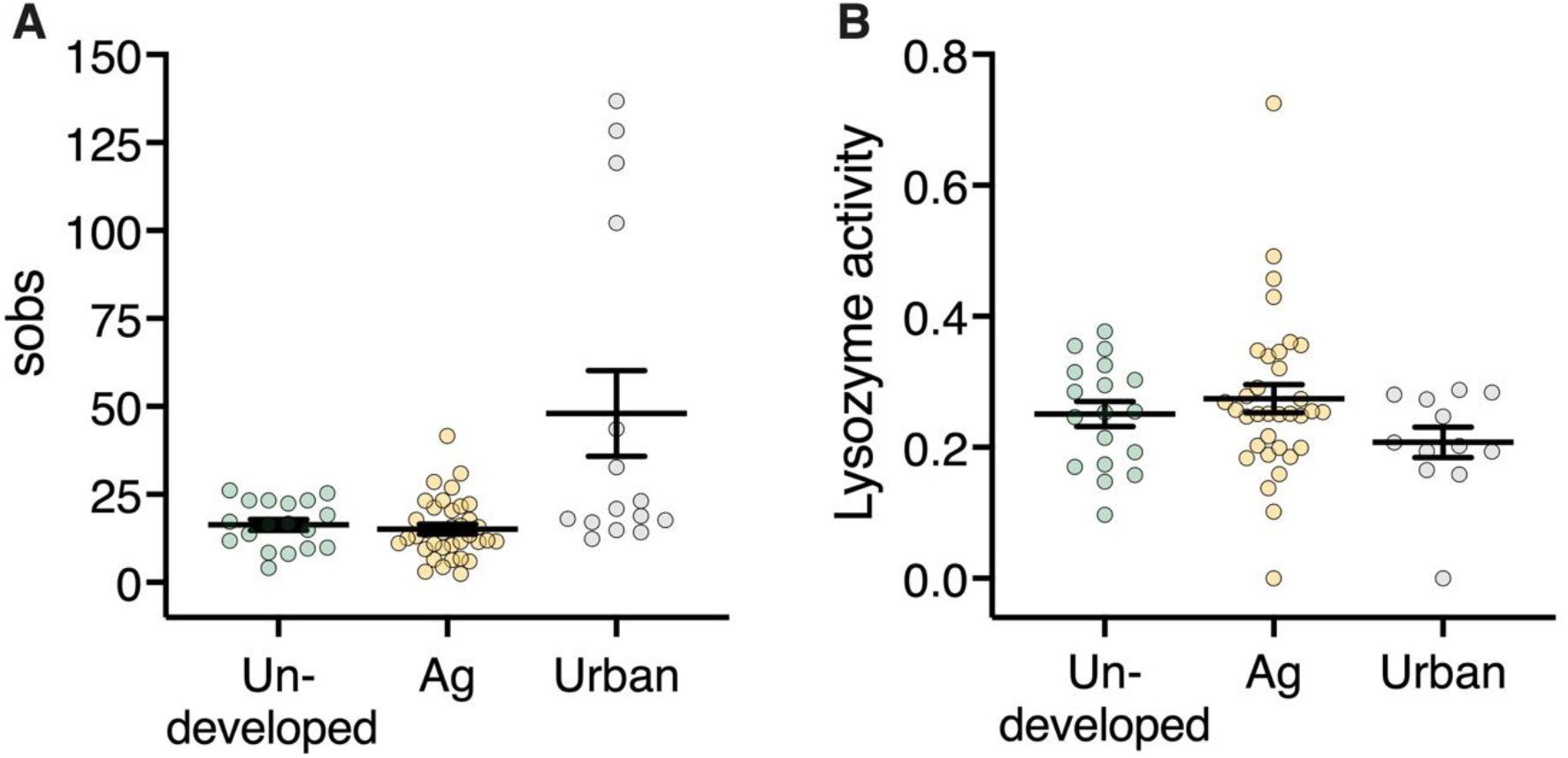
The effect of land use type (undeveloped, agricultural [Ag], urban) on mean (± SE) sobs bacterial diversity (A) and lysozyme activity (B). Each point represents an individual.

Bacterial community structure (*F* = 1.33, *P* = 0.04) and membership (*F* = 1.19, *P* = 0.045) differed between urban areas compared to the other land use types for medium ground finches (Fig. 2A-B). However, bacterial community structure (*F* = 0.71, *P* = 0.98) and membership (*F* = 0.83, *P* = 0.98) did not differ significantly among land uses in small ground finches (Fig. 2A-B). For small ground finches, relative abundances of several genera, including genera *Pseudoxanthomonas, Cloacibacterium*, and *Dietzia*, were higher in urban areas compared to the other two areas (Table S3). Relative abundances of phyla did not differ among land use types for small or medium ground finches (*P* > 0.05). Relative abundances of genera did not differ among land use types for medium ground finches either (*P* > 0.05).

**Fig. 2.**
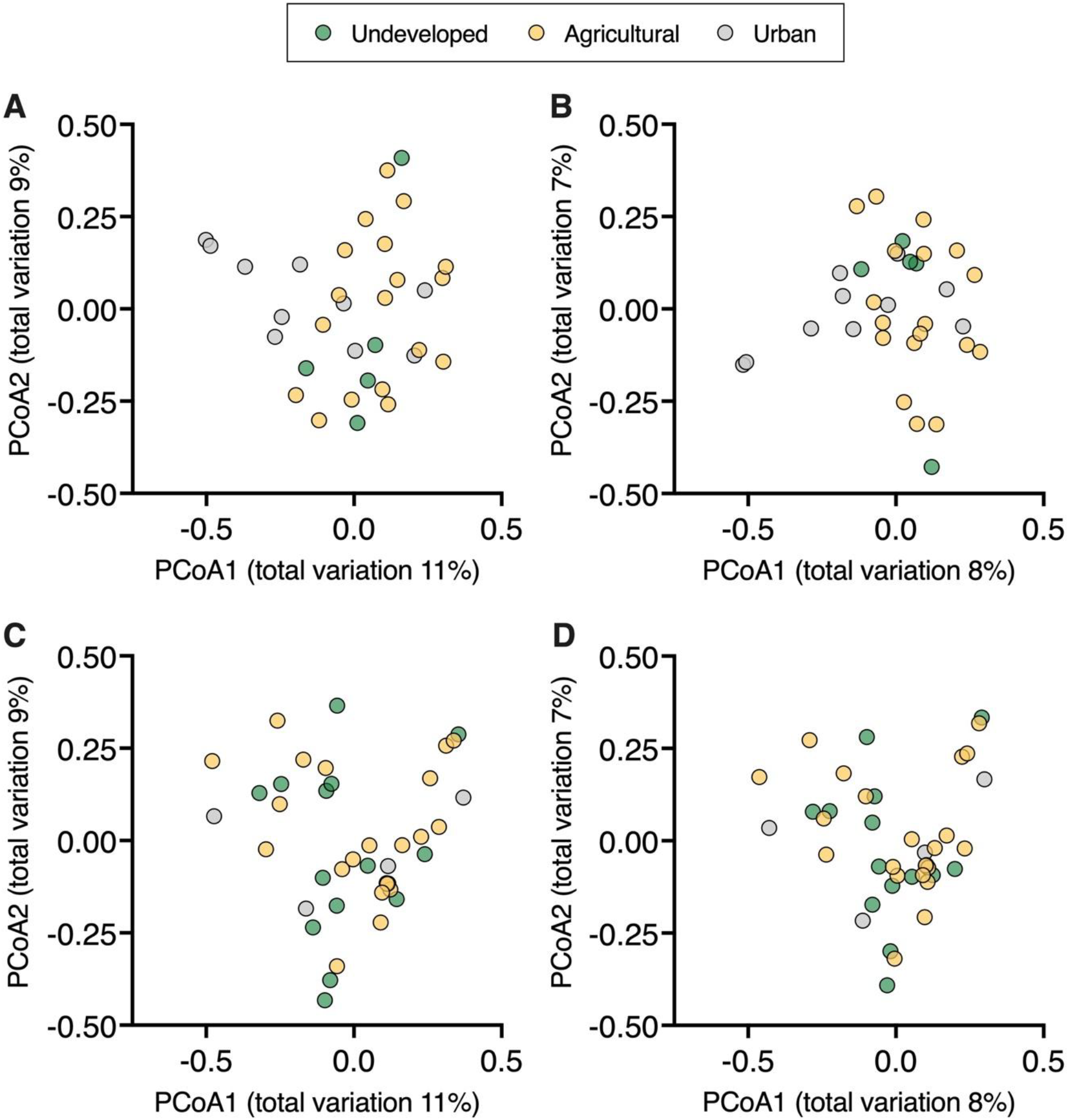
The effect of land use type on bacterial community structure and membership for medium ground finches (A: structure, B: membership) and small ground finches (C: structure, D: membership).

### Effect of land use and finch species on the immune response

Lysozyme activity differed among land use types for the medium and small ground finches (Fig. 1B; Table S4). Overall, urban finches had statistically lower lysozyme levels than finches in the agricultural and undeveloped areas, but this effect was more pronounced in medium ground finches (Fig. 1B; Table S5). Haptoglobin, complement antibody, and natural antibody levels did not differ significantly among land use types in either medium or small ground finches (Table S4-S5). Immune metrics did not correlate significantly with the bacterial diversity metrics (R^2^ < 0.20 for all pairs).

### Effect of land use and finch species on body size and mass

Body mass differed between small and medium ground finches (Fig. 3; Table S6-S7). For small ground finches, body mass did not differ significantly among land use types, but for medium ground finches, individuals in the undeveloped area had lower body mass than individuals in agricultural and urban areas (Fig. 3; Table S7). Tarsus length and scaled mass index differed significantly between species but not among land use types (Table S6-S7).

**Fig. 3.**
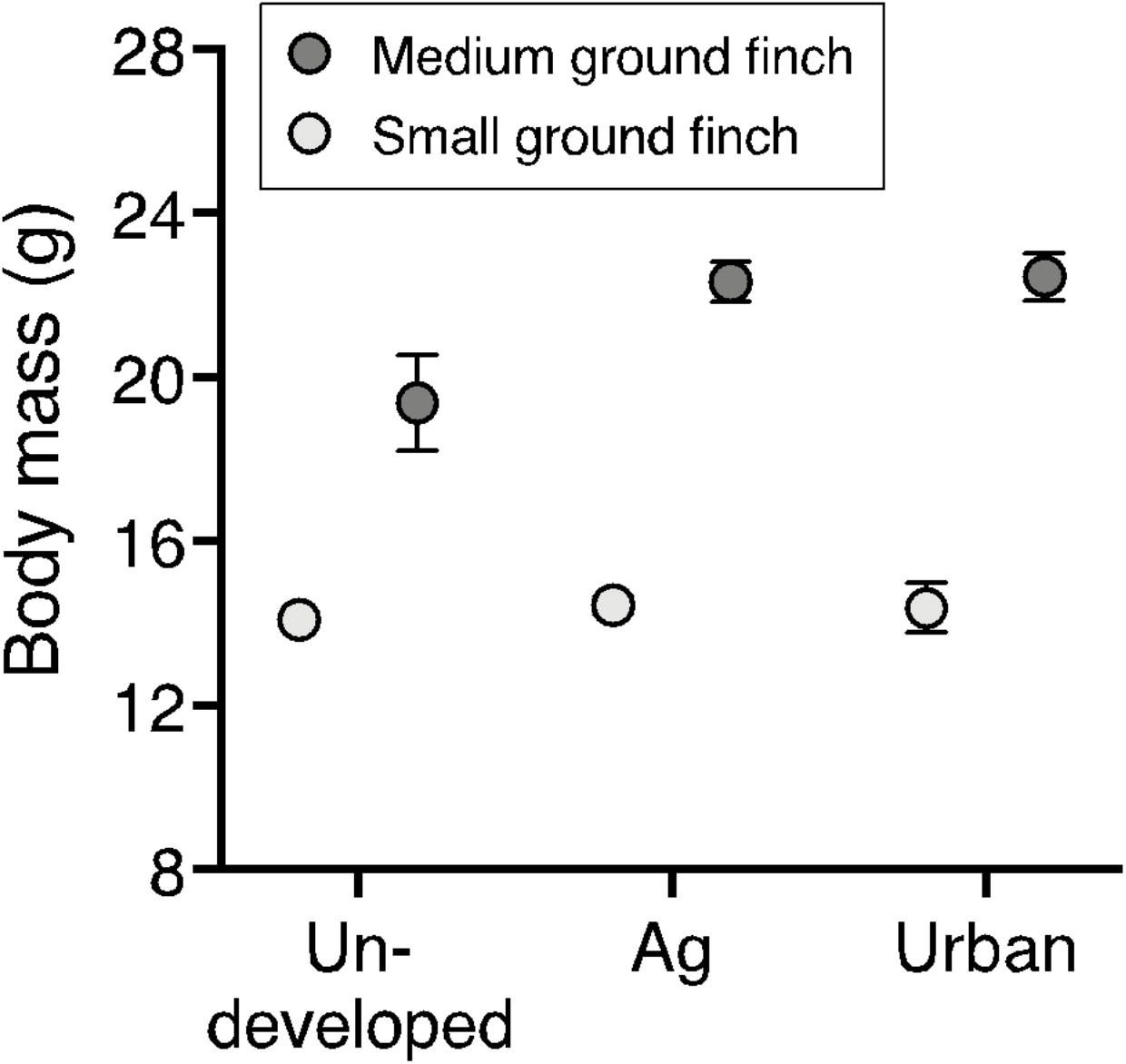
The effect of land use type (undeveloped, agricultural [Ag], urban) and finch species (small ground finch [SGF], and medium ground finch [MGF]) on body mass.

## DISCUSSION

Our study examined the effect of land use on the gut microbiota, immune metrics, and body measurements of two Darwin’s finch species in 2008, at the start of a rapid increase in human activity in the Galápagos Islands. Studies of samples collected in 2016 have found significant impacts of human activity on metrics of the gut microbiota and body measurements of Darwin’s finches. Based on data collected eight years earlier, we found marginal effects of urbanization on the observed species richness (sobs) across bird species and a few bacterial genera (*Pseudoxanthomonas, Cloacibacterium*, and *Dietzia*) in small ground finches. Similarly, we found few effects of land use on immune metrics and body measurements, with urbanization affecting lysozyme levels across bird species and body mass in medium ground finches. Furthermore, bacterial diversity did not correlate with any of the immune measures. Although humans have had a permanent presence in the Galápagos for decades, our results suggest that initially, increased human activity (starting ∼2007) had marginal effects on the finches.

Lysozyme activity, but not the other immune metrics, was influenced by land use type; urban birds had lower lysozyme activity than non-urban birds. Lysozymes are part of the innate immune system, which are transferred from mother to egg whites. During the breeding season, lysozyme activity declines in females during pre-laying and egg laying (Saino *et al*. 2002). Urban Darwin’s finches differ in the length and timing of their breeding season compared to non-urban birds (Harvey *et al*. 2021). Samples were collected from female finches in February, which is early in the breeding season. Therefore, timing of breeding could be responsible for differences in lysozyme activity in urban vs. non-urban birds. Furthermore, although lysozyme is an antimicrobial enzyme, we found no relationship between lysozyme activity and bacterial diversity. This lack of relationship is likely because the lysozymes quantified in our study were in the plasma, rather in the gut mucosal tissue. Overall, because lysozyme activity differs across land uses in 2008, early in urbanization, future studies could focus on how lysozyme activity has changed in finches since then, especially in response to emerging diseases (Wikelski *et al*. 2004).

Observed species richness (sobs) was higher in urban areas compared to non-urban areas. However, the other alpha diversity metrics that consider species evenness did not differ across habitats. Other studies that have found higher observed species richness and Shannon index in urban birds indicate that this could be caused by man-made landscapes and a higher grass and tree cover (Phillips *et al*., 2018; Berlow *et al*., 2021). However, there have also been studies demonstrating less bacterial diversity in species living in an urban habitat when compared to species living in a non-urban habitat (Barelli *et al*., 2015; Teyssier *et al*., 2018, 2020; Knutie *et al*., 2019). A less diverse gut community in urban areas could be caused by habitat disturbances or differences in diets between urban and non-urban populations (Sonnenburg *et al*., 2016; Teyssier *et al*., 2018, 2020; Knutie, 2020). The time of sampling should also be considered, as Tessyier *et al*. (2018) found that birds in a non-urban habitat exhibited less microbiome diversity during the winter months, while microbiome diversity in birds in an urban habitat did not differ across seasons. As the birds included in this study were sampled at the start of the rainy/breeding season, it is possible that a seasonal effect contributed to the difference seen between birds from non-urban and urban habitats. Overall, our results contribute to the growing body of literature demonstrating that there are many factors to consider when studying the relationship between urbanization and gut microbiome diversity.

Relative abundances of several genera, including *Pseudoxanthomonas* (phylum Proteobacteria), *Cloacibacterium* (phylum Bacteroidetes), and *Dietzia* (phylum Actinobacteria), were higher in urban areas than non-urban areas in small ground finches in 2008. Studies have not specifically documented differences in these genera across land use types but have found higher relative abundances of family Weeksellaceae (includes *Cloacibacterium spp*.) in urban areas compared to non-urban areas (Berlow *et al*., 2021). Interestingly, small ground finches in 2016 only showed increases in the relative abundance of *Steroidobacter spp*. in the presence of humans (Knutie *et al*., 2019). Although we found no effect of human activity on bacterial phyla in 2008 finches, 2016 finches in areas with higher human activity had higher relative abundances of phyla Chlamydiae and Cyanobacteria than in other areas.

One explanation for differences in bacterial taxa between 2008 and 2016 finches is that the gut microbiota of finches has changed over eight years coinciding with a change in the finches’ environment. Davidson et al., (2020) found that relative abundances of phyla Proteobacteria, Bacteroidetes, Actinobacteria in great tits (*Parus major*) were higher in urban areas compared to rural areas and/or when tits were fed an insect-based diet compared to a seed-based diet. Recent studies have found that urban small and medium ground finches tend to prefer human food diets (e.g., chips and cooked rice) rather than their natural diet (e.g., seeds) which could explain our results (de León *et al*., 2018). Alternatively, urban contaminants could be responsible for changes in the gut bacterial taxa observed in the finches. *Pseudoxanthomonas spp*. have been found in oil-polluted water bodies and soil because the bacterial taxa can metabolize and degrade the contaminant (Young *et al*., 2007). Although human activity in the Islands did not rapidly increase until approximately 2007, sources of human-driven pollution were present in towns such as Puerto Ayora. For example, the major presence of vehicles began in the 2000s with the completion of major paved roads. Although contaminants can affect the abundance of bacterial taxa, medium ground finches were likely exposed to the same environmental contaminants as the small ground finches but, nonetheless, did not show differences in bacterial taxa across land uses. However, since small and medium ground finches do eat different natural diets, perhaps the natural diet of the small ground finches shifted in response to urbanization over time, while the medium ground finches did not.

Body mass differed across land-use types for medium ground finches, with non-urban birds having lower body mass than those in urban areas. These results were also found by McNew *et al*. (2017) and can potentially be explained by differences in food availability across sites; human activity can affect food availability and preference for wildlife. For example, the diet of non-urban birds includes mostly natural foods, such as seeds, fruits, and insects, while urban birds can prefer human processed food (Murray *et al*., 2016; de León *et al*., 2018; Phillips *et al*., 2018). A human-based diet can affect body morphometrics of animals, including increased body mass (Banks and Dickman, 2000; Bayol *et al*., 2007; Wilcoxen *et al*., 2015). In contrast, we found no effect of land use on the body mass of small ground finches. Small ground finches have smaller bill sizes (length, width, depth) and therefore different diets than medium ground finches (Abbott *et al*., 1977). One explanation is that perhaps medium ground finches are better able to exploit the urban diet than small ground finches. Interestingly, small ground finches in areas with more human activity in 2016 were larger than in areas with no human activity, which might be related to the increase in human activity over time. Since museum specimens and long-term field data exist, a future study could determine the effect of human activity on different finch species in urban and non-urban areas across islands.

## CONCLUSION

Our study suggests that immunity, gut microbiota, and body mass of two species of Darwin’s finches vary across land use prior to the rapid increase in human activity in Galápagos Islands. These results suggest that earlier human activity can affect the ecology of birds. Over the past several decades, Darwin’s finches have faced increasing challenges from invasive parasites (Wikelski *et al*., 2004; B. Fessl *et al*., 2010; Koop *et al*., 2016; Knutie, 2018) and predators (Gotanda, 2021), anthropogenic debris (Theodosopoulos and Gotanda, 2018; Harvey *et al*., 2021), and dynamic annual changes in natural and novel food availability (Grant and Grant, 1995; de León *et al*., 2018), which can all affect the physiology and gut microbiota of animals. Given our results and the novel challenges facing the Galápagos Islands, the iconic Darwin’s finch system has exciting potential for future ecoimmunology and microbiome research in a changing world (Ohmer *et al*., 2021).

## Supporting information

Supplemental Results

## ACKNOWLEDGEMENT

We thank Kelly Lee and Rachel Mills for field assistance; Martin Wikelski and Patricia Parker for logistic support; Kirk Klasing for use of laboratory facilities; the Galápagos National Park (GNP) and the Charles Darwin Research station for logistical support; and Kendra Maas at the University of Connecticut Microbial Analysis, Resources, and Services for workshop training on the Mothur bioinformatics. The work was supported by funding through American Ornithologists’ Union, Sigma Xi, University of California, Davis Office of Graduate Studies, a National Science Foundation Graduate Research Fellowship, and TAME (providing discount airfare) to MZ, and start-up funds and a Research Excellence Program Grant from the University of Connecticut to SAK. Our work was done under GNP permits #0499-2008-PNG/DIR. All applicable international, national, and/or institutional guidelines for the care and use of animals were followed (UC Davis IACUC protocol 13171).

## Authors’ contributions

MZ and SAK conceived the study, MZ collected the samples and field data, MZ and AA conducted the laboratory work, AA conducted the bioinformatics, JNB, AA, and SAK did the data analyses. All authors wrote, revised, and approved the manuscript.

## Data accessibility

Data are available at FigShare (doi: available upon acceptance) and sequences have been uploaded to GenBank (BioProject accession number: available upon acceptance).

## LITERATURE CITED

Abbott I, Abbott LK, Grant PR (1977) Comparative Ecology of Galápagos Ground Finches (Geospiza Gould): Evaluation of the Importance of Floristic Diversity and Interspecific Competition. Ecological Monographs 47. doi:10.2307/1942615

Atwell JW, Cardoso GC, Whittaker DJ, Campbell-Nelson S, Robertson KW, Ketterson ED (2012) Boldness behavior and stress physiology in a novel urban environment suggest rapid correlated evolutionary adaptation. Behavioral Ecology 23: 960–969.

Banks PB, Dickman CR (2000) Effects of winter food supplementation on reproduction, body mass, and numbers of small mammals in montane Australia. Canadian Journal of Zoology 78. doi:10.1139/z00-110

Barelli C, Albanese D, Donati C, Pindo M, Dallago C, Rovero F, Cavalieri D, Michael Tuohy K, Christine Hauffe H, de Filippo C (2015) Habitat fragmentation is associated to gut microbiota diversity of an endangered primate: Implications for conservation. Scientific Reports 5. doi:10.1038/srep14862

Bates D, Mächler M, Bolker BM, Walker SC (2015) Fitting linear mixed-effects models using lme4. Journal of Statistical Software. doi:10.18637/jss.v067.i01

Bayol SA, Farrington SJ, Stickland NC (2007) A maternal “junk food” diet in pregnancy and lactation promotes an exacerbated taste for “junk food” and a greater propensity for obesity in rat offspring. British Journal of Nutrition 98: 843–51.

Berlow M, Phillips JN, Derryberry EP (2021) Effects of Urbanization and Landscape on Gut Microbiomes in White-Crowned Sparrows. Microbial Ecology 81. doi:10.1007/s00248-020-01569-8

Bosse M, Spurgin LG, Laine VN, Cole EF, Firth JA, Gienapp P, Gosler AG, McMahon K, Poissant J, Verhagen I, et al. (2017) Recent natural selection causes adaptive evolution of an avian polygenic trait. Science 6361: 365–368.

Caporaso JG, Lauber CL, Walters WA, Berg-Lyons D, Huntley J, Fierer N (2012) Ultra-high-throughput microbial community analysis on the Illumina HiSeq and MiSeq platforms. ISME J 6. doi:10.1038/ismej.2012.8

Caroll MC, Prodeus AP (1998) Linkages of innate and adaptive immunity. Current Opinion in Immunology 10. doi:10.1016/S0952-7915(98)80028-9

Casali P, Schettino EW (1996) Structure and function of natural antibodies. Current Topics in Microbiology and Immunology 210. doi:10.1007/978-3-642-85226-8_17

Cornet S, Bichet C, Larcombe S, Faivre B, Sorci G (2014) Impact of host nutritional status on infection dynamics and parasite virulence in a bird-malaria system. The Journal of Animal Ecology 83: 256–265.

Davidson GL, Wiley N, Cooke AC, Johnson CN, Fouhy F, Reichert MS, de la Hera I, Crane JMS, Kulahci IG, Ross RP, et al. (2020) Diet induces parallel changes to the gut microbiota and problem solving performance in a wild bird. Scientific Reports 10. doi:10.1038/s41598-020-77256-y

de León L, Sharpe D, Gotanda K, Raeymaekers J, Chaves J, Hendry A, Podos J (2018) Urbanization erodes niche segregation in Darwin’s finches. Evolutionary Applications.

Demas GE (2004) The energetics of immunity: a neuroendocrine link between energy balance and immune function. Hormones and Behavior.

Edgar RC, Haas BJ, Clemente JC, Quince C, Knight R (2011) UCHIME improves sensitivity and speed of chimera detection. Bioinformatics 27: 2194–2200.

Fessl Birgit, Young GH, Young RP, Rodríguez-Matamoros J, Dvorak M, Tebbich S, Fa JE (2010) How to save the rarest Darwin’s finch from extinction: the mangrove finch on Isabela Island. Philosophical Transactions of the Royal Society of London B 365: 1019– 1030.

Fessl B., Young GH, Young RP, Rodríguez-Matamoros J, Dvorak M, Tebbich S, Fa JE (2010) How to save the rarest Darwin’s finch from extinction: the mangrove finch on Isabela Island. Philosophical Transactions of the Royal Society of London B 365: 1019–1030.

Fox J, Weisberg S (2019) CAR - An R Companion to Applied Regression. Thousand Oaks CA: Sage. Gotanda KM (n.d.) Human influences on antipredator behaviour in Darwin’s finches.

Grant PR, Grant BR (1995) Predicting microevolutionary responses to directional selection on heritable variation. Evolution 49: 241–251.

Grond K, Sandercock BK, Jumpponen A, Zeglin LH (2018) The avian gut microbiota: community, physiology and function in wild birds. Journal of Avian Biology 49: e01788.

Harvey JA, Chernicky K, Simons SR, Verrett TB, Chaves JA, Knutie SA (2021) Urban living influences the nesting success of Darwin’s finches in the Galápagos Islands. Ecology and Evolution 11. doi:10.1002/ece3.7360

Hendry AP, Grant PR, Rosemary Grant B, Ford HA, Brewer MJ, Podos J (2006) Possible human impacts on adaptive radiation: beak size bimodality in Darwin’s finches. Proceedings of the Royal Society B: Biological Sciences 273: 1887–94.

Hirsch RL (1982) The complement system: Its importance in the host response to viral infection. Microbiological Reviews.

Howick VM, Lazzaro BP (2014) Genotype and diet shape resistance and tolerance across distinct phases of bacterial infection. BMC Evolutionary Biology 14: 56.

Jin Y, Wu S, Zeng Z, Fu Z (2017) Effects of environmental pollutants on gut microbiota. Environmental Pollution.

Knutie SA (2018) Relationships among introduced parasites, host defenses, and gut microbiota of Galápagos birds. Ecosphere 9: e02286.

Knutie SA (2020) Food supplementation affects gut microbiota and immunological resistance to parasites in a wild bird species. Journal of Applied Ecology 57. doi:10.1111/1365-2664.13567

Knutie SA, Chaves JA, Gotanda KM (2019) Human activity can influence the gut microbiota of Darwin’s finches in the Galápagos Islands. Molecular Ecology mec.15088.

Koop JAH, Kim PS, Knutie SA, Adler F, Clayton DH (2016) An introduced parasitic fly may lead to local extinction of Darwin’s finch populations. Journal of Applied Ecology 53. doi:10.1111/1365-2664.12575

Kozich JJ, Westcott SL, Baxter NT, Highlander SK, Schloss PD (2013) Development of a dual-index sequencing strategy and curation pipeline for analyzing amplicon sequence data on the miseq illumina sequencing platform. Applied and Environmental Microbiology 79: 5112–5120.

Kumar PS, Mason MR, Brooker MR, O’Brien K (2012) Pyrosequencing reveals unique microbial signatures associated with healthy and failing dental implants. Journal of Clinical Periodontology 39: 425–433.

Lochmiller RL, Deerenberg C (2000) Trade-Offs in Evolutionary Immunology: just what Is the cost of immunity? Oikos 88: 87–98.

Loo WT, García-Loor J, Dudaniec RY, Kleindorfer S, Cavanaugh CM (2019) Host phylogeny, diet, and habitat differentiate the gut microbiomes of Darwin’s finches on Santa Cruz Island. Scientific Reports 9. doi:10.1038/s41598-019-54869-6

Matson KD, Ricklefs RE, Klasing KC (2005) A hemolysis-hemagglutination assay for characterizing constitutive innate humoral immunity in wild and domestic birds. Developmental and Comparative Immunology 29. doi:10.1016/j.dci.2004.07.006

McNew SM, Beck D, Sadler-Riggleman I, Knutie SA, Koop JAH, Clayton DH, Skinner MK (2017) Epigenetic variation between urban and rural populations of Darwin’s finches. BMC Evolutionary Biology 17. doi:10.1186/s12862-017-1025-9

Michel AJ, Ward LM, Goffredi SK, Dawson KS, Baldassarre DT, Brenner A, Gotanda KM, McCormack JE, Mullin SW, O’Neill A, et al. (2018) The gut of the finch: uniqueness of the gut microbiome of the Galápagos vampire finch. Microbiome 6: 167.

Millet S, Bennett J, Lee KA, Hau M, Klasing KC (2007) Quantifying and comparing constitutive immunity across avian species. Developmental & Comparative Immunology 31: 188–201.

Møller AP (2008) Flight distance of urban birds, predation, and selection for urban life. Behavioral Ecology and Sociobiology 63: 63.

Murphy K, Weaver C, Janeway C (2017) Janeway ‘s Immunobiology 9Th Edition. America.

Murray MH, Becker DJ, Hall RJ, Hernandez SM (2016) Wildlife health and supplemental feeding: A review and management recommendations. Biological Conservation 204: 163– 174.

Ohmer MEB, Costantini D, Czirják G, Downs CJ, Ferguson L v., Flies A, Franklin CE, Kayigwe AN, Knutie S, Richards-Zawacki CL, et al. (2021) Applied ecoimmunology: Using immunological tools to improve conservation efforts in a changing world. Conservation Physiology.

Oksanen J, Blanchet FG, Friendly M, Kindt R, Legendre P, Mcglinn D, Minchin PR, O’Hara RB, Simpson GL, Solymos P, et al. (2019) vegan: Community Ecology Package. R package version 2.4-2. Community ecology package 2.5-6.

Peig J, Green AJ (2009) New perspectives for estimating body condition from mass/length data: the scaled mass index as an alternative method. Oikos 118: 1883–1891.

Phillips JN, Berlow M, Derryberry EP (2018) The effects of landscape urbanization on the gut microbiome: an exploration into the gut of urban and rural white-crowned sparrows. Frontiers in Ecology and Evolution 6: 1–10.

Quast C, Pruesse E, Yilmaz P, Gerken J, Schweer T, Yarza P, Peplies J, Glöckner FO (2013) The SILVA ribosomal RNA gene database project: improved data processing and web-based tools. Nucleic Acids Research 41: D590–D596.

Quaye IK (2008) Haptoglobin, inflammation and disease. Transactions of the Royal Society of Tropical Medicine and Hygiene.

Round June L, Mazmanian SK (2009) The gut microbiota shapes intestinal immune responses during health and disease. Nature Reviews Immunology 9: 313–323.

Round June L., Mazmanian SK (2009) The gut microbiota shapes intestinal immune responses during health and disease. Nature Reviews Immunology 9: 313–323.

Samia DSM, Blumstein DT, Díaz M, Grim T, Ibáñez-Álamo JD, Jokimäki J, Tätte K, Markó G, Tryjanowski P, Møller AP (2017) Rural-Urban Differences in Escape Behavior of European Birds across a Latitudinal Gradient. Frontiers in Ecology and Evolution 5: 1–13.

Schloss PD, Westcott SL, Ryabin T, Hall JR, Hartmann M, Hollister EB, Lesniewski RA, Oakley BB, Parks DH, Robinson CJ, et al. (2009) Introducing mothur: Open-source, platform-independent, community-supported software for describing and comparing microbial communities. Applied and Environmental Microbiology 75: 7537–7541.

Shchipkova AY, Nagaraja HN, Kumar PS (2010) Subgingival microbial profiles of smokers with periodontitis. Journal of Dental Research 89: 1247–1253.

Sheldon BC, Verhulst S (1996) Ecological immunology: costly parasite defences and trade-offs in evolutionary ecology. Trends in Ecology & Evolution 11: 317–321.

Sol D, Maspons J, Gonzalez-Voyer A, Morales-Castilla I, Garamszegi LZ, Møller AP (2018) Risk-taking behavior, urbanization and the pace of life in birds. Behavioral Ecology and Sociobiology 72: 58–66.

Sonnenburg ED, Smits SA, Tikhonov M, Higginbottom SK, Wingreen NS, Sonnenburg JL (2016) Diet-induced extinctions in the gut microbiota compound over generations. Nature 529. doi:10.1038/nature16504

Start D, Bonner C, Weis AE, Gilbert B (2018) Consumer-resource interactions along urbanization gradients drive natural selection. Evolution 9: 1863–1873.

Sternberg ED, Lefèvre T, Li J, de Castillejo CLF, Li H, Hunter MD, de Roode JC (2012) Food plant derived disease tolerance and resistance in a natural butterfly-plant-parasite interactions. Evolution 66: 3367–3376.

Strandin T, Babayan SA, Forbes KM (2018) Reviewing the effects of food provisioning on wildlife immunity. Philosophical Transactions of the Royal Society B: Biological Sciences 373: 20170088.

Svensson E, Råberg L, Koch C, Hasselquist D (1998) Energetic stress, immunosuppression and the costs of an antibody response. Functional Ecology 12: 912–919.

Teyssier A, Matthysen E, Hudin NS, de Neve L, White J, Lens L (2020) Diet contributes to urban-induced alterations in gut microbiota: Experimental evidence from a wild passerine. Proceedings of the Royal Society B: Biological Sciences 287. doi:10.1098/rspb.2019.2182

Teyssier A, Rouffaer LO, Saleh Hudin N, Strubbe D, Matthysen E, Lens L, White J (2018) Inside the guts of the city: Urban-induced alterations of the gut microbiota in a wild passerine. Science of the Total Environment 612: 1276–1286.

Theodosopoulos AN, Gotanda KM (2018) Death of a Darwin’s Finch: a consequence of human-made debris? The Wilson Journal of Ornithology. doi:10.1676/17-00050.1

Walsh SJ, Mena CF (2016) Interactions of social, terrestrial, and marine sub-systems in the Galápagos Islands, Ecuador. Proceedings of the National Academy of Sciences 113: 14536– 14543.

Wang Q, Garrity GM, Tiedje JM, Cole JR (2007) Naïve Bayesian classifier for rapid assignment of rRNA sequences into the new bacterial taxonomy. Applied and Environmental Microbiology 73: 5261–5267.

Watkins G, Cruz F (2007) Galápagos at Risk: A Socioeconomic Analysis of the Situation in the Archipelago. Puerto Ayora, Santa Cruz.

Wicher KB, Fries E (2006) Haptoglobin, a hemoglobin-binding plasma protein, is present in bony fish and mammals but not in frog and chicken. Proceedings of the National Academy of Sciences of the United States of America 103. doi:10.1073/pnas.0508723103

Wickham H (2007) Reshaping data with the reshape package. Journal of Statistical Software 21. doi:10.18637/jss.v021.i12

Wickham H (2011) The split-apply-combine strategy for data analysis. Journal of Statistical Software 40. doi:10.18637/jss.v040.i01

Wickham H, Henry L (2019) tidyr: Tidy Messy Data. R package version 100.

Wikelski M, Foufopoulos J, Vargas H, Snell H (2004) Galápagos birds and diseases: invasive pathogens as threats for island species. Ecology and Society 9: online: http://www.ecologyandsociety.org/vol9/iss1.

Wilcoxen TE, Horn D, Hogan BM, Hubble CN, Huber SJ, Flamm J, Knott M, Lundstrom L, Salik F, Wassenhove SJ, et al. (2015) Effects of bird-feeding activities on the health of wild birds. Conservation Physiology 3: cov058.

Young CC, Ho MJ, Arun AB, Chen WM, Lai WA, Shen FT, Rekha PD, Yassin AF (2007) Pseudoxanthomonas spadix sp. nov., isolated from oil-contaminated soil. International Journal of Systematic and Evolutionary Microbiology 57. doi:10.1099/ijs.0.65053-0

Zylberberg M, Klasing KC, Hahn TP (2014) In house finches, Haemorhous mexicanus, risk takers invest more in innate immune function. Animal Behaviour 89. doi:10.1016/j.anbehav.2013.12.021

Zylberberg M, Lee KA, Klasing KC, Wikelski M (2013) Variation with land use of immune function and prevalence of avian pox in Galápagos finches. Conservation Biology 27: 103–112.

