## Supplemental Results for "Influence of human activity on gut microbiota and immune responses of Darwin’s finches in the Galápagos Islands"

Supplemental Table 1. The results of GLMs on the effect of land use type, finch species, and their interaction on bacterial diversity metrics.

|  | Land use type | Species | Interaction |
| --- | --- | --- | --- |
| sobs | χ^2^ = 16.76, df = 2,  ***P* = 0.0002** | χ^2^ = 1.66, df = 1,  *P* = 0.20 | χ^2^ = 2.60, df = 2,  *P* = 0.27 |
| Shannon index | χ^2^ = 4.35, df = 2,  *P* = 0.11 | χ^2^ =2.53, df = 1,  *P* = 0.11 | χ^2^ =2.04, df = 2,  *P* = 0.36 |
| Simpson index | χ^2^ = 2.28, df = 2,  *P* = 0.32 | χ^2^ = 1.52, df = 1,  *P* = 0.22 | χ^2^ = 2.03, df = 2,  *P* = 0.36 |
| Shannon Evenness | χ^2^ = 1.04, df = 2,  *P* = 0.60 | χ^2^ = 1.66, df = 1,  *P* = 0.20 | χ^2^ = 0.66, df = 2,  *P* = 0.72 |

Supplemental Table 2. Mean (±SE) of each bacterial diversity metric across land use types and finch species. Sample size in parenthesis.

|  | Undeveloped | Agricultural | Urban |
| --- | --- | --- | --- |
| Small ground finches |  |  |  |
| sobs | 14.91 ± 1.88 (13) | 17.15 ± 2.22 (18) | 46.96 ± 23.0 (5) |
| Shannon index | 1.76 ± 0.17 (13) | 1.82 ± 0.14 (18) | 2.19 ± 0.51 (5) |
| Simpson index | 5.82 ± 0.74 (13) | 5.66 ± 0.65 (18) | 9.86 ± 5.10 (5) |
| Shannon Evenness | 0.67 ± 0.04 (13) | 0.66 ± 0.03 (18) | 0.63 ± 0.09 (5) |
| Medium ground finches |  |  |  |
| sobs | 20.01 ± 2.45 (5) | 13.12 ± 1.65 (18) | 48.53 ± 15.08 (10) |
| Shannon index | 1.98 ± 0.20 (5) | 1.52 ± 0.14 (18) | 1.92 ± 0.13 (10) |
| Simpson index | 6.58 ± 1.19 (5) | 4.47 ± 0.54 (18) | 4.79 ± 0.77 (10) |
| Shannon Evenness | 0.66 ± 0.04 (5) | 0.61 ± 0.04 (18) | 0.57 ± 0.05 (10) |

Supplemental Table 3. Mean (±SE) relative abundances (%) of bacterial genera across land use types for small ground finches. Sample size in parenthesis.

| Genus | Undeveloped | Agricultural | Urban | Statistics |
| --- | --- | --- | --- | --- |
| *Pseudoxanthomonas* | 0.00 ± 0.00% (14) | 0.00 ± 0.00% (21) | 0.38 ± 0.23% (5) | *F* = 10.74, ***P* = 0.01** |
| *Cloacibacterium* | 0.00 ± 0.00% (14) | 0.00 ± 0.00% (21) | 0.38 ± 0.23% (5) | *F* = 10.74, ***P* = 0.01** |
| *Dietzia* | 0.22 ± 0.22% (14) | 0.00 ± 0.00% (21) | 1.25 ± 0.80% (5) | *F* = 8.35, ***P* = 0.04** |

Supplemental Table 4. The results of GLMs on the effect of land use type, finch species, and their interaction on immune metrics.

|  | Land use type | Species | Interaction |
| --- | --- | --- | --- |
| Haptoglobin | χ^2^ = 0.77, df = 2,  *P* = 0.68 | χ^2^ = 0.26, df = 1,  *P* = 0.61 | χ^2^ = 2.25, df = 2,  *P* = 0.33 |
| Lysozyme | χ^2^ = 6.86, df = 2,  ***P* = 0.03** | χ^2^ = 1.82, df = 1,  *P* = 0.18 | χ^2^ = 2.42, df = 2,  *P* = 0.30 |
| Complement antibodies | χ^2^ = 2.85, df = 2,  *P* = 0.24 | χ^2^ = 0.63, df = 1,  *P* = 0.43 | χ^2^ = 0.55, df = 2,  *P* = 0.76 |
| Natural antibodies | χ^2^ = 4.14, df = 2,  *P* = 0.13 | χ^2^ = 1.72, df = 1,  *P* = 0.19 | χ^2^ = 3.61, df = 2,  *P* = 0.16 |

Supplemental Table 5. Mean (±SE) of each immune metric across land use types and finch species. Sample size in parenthesis.

|  | Undeveloped | Agricultural | Urban |
| --- | --- | --- | --- |
| Small ground finches |  |  |  |
| Haptoglobin | 0.97 ± 0.16 (13) | 0.94 ± 0.08 (20) | 0.52 ± 0.14 (3) |
| Lysozyme | 0.23 ± 0.02 (13) | 0.25 ± 0.02 (21) | 0.24 ± 0.03 (3) |
| Complement antibodies | 3.17 ± 0.55 (12) | 5.00 ± 0.48 (20) | 5.50 ± 0.50 (2) |
| Natural antibodies | 1.21 ± 0.28 (12) | 1.00 ± 0.18 (20) | 1.50 ±1.0 (2) |
| Medium ground finches |  |  |  |
| Haptoglobin | 0.81 ± 0.18 (5) | 0.89 ± 0.09 (18) | 0.74 ± 0.07 (10) |
| Lysozyme | 0.31 ± 0.02 (5) | 0.31 ± 0.05 (14) | 0.20 ± 0.03 (9) |
| Complement antibodies | 3.40 ± 0.51 (5) | 5.53 ± 0.53 (17) | 5.10 ± 0.74 (10) |
| Natural antibodies | 0.60 ± 0.29 (5) | 1.32 ± 0.19 (17) | 1.50 ± 0.37 (10) |

Supplemental Table 6. The results of GLMs on the effect of land use type, finch species, and their interaction on body mass and size.

|  | Land use type | Species | Interaction |
| --- | --- | --- | --- |
| Body mass | χ^2^ = 12.08, df = 2,  ***P* = 0.002** | χ^2^ = 237.52, df = 1,  ***P* < 0.0001** | χ^2^ = 5.70, df = 2,  ***P* = 0.058** |
| Tarsus length | χ^2^ = 5.20, df = 2,  *P* = 0.07 | χ^2^ = 60.98, df = 1,  ***P* < 0.0001** | χ^2^ = 2.15, df = 2,  *P* = 0.34 |
| Scaled mass index | χ^2^ = 1.22, df = 2,  *P* = 0.54 | χ^2^ = 233.02, df = 1,  ***P* < 0.0001** | χ^2^ = 0.50, df = 2,  *P* = 0.78 |

Supplemental Table 7. Mean (±SE) of each body mass (g) and tarsus length (mm) across land use types and finch species. Sample size in parenthesis.

|  | Undeveloped | Agricultural | Urban |
| --- | --- | --- | --- |
| Small ground finches |  |  |  |
| Body mass | 14.07 ± 0.30 (14) | 14.43 ± 0.23 (21) | 14.36 ± 0.61 (5) |
| Tarsus length | 21.12 ± 0.31 (14) | 21.41 ± 0.22 (21) | 21.30 ± 0.34 (5) |
| Scaled mass index | 14.27 ± 0.26 (14) | 14.33 ± 0.23 (21) | 14.37 ± 0.62 (5) |
| Medium ground finches |  |  |  |
| Body mass | 19.36 ± 1.17 (5) | 22.32 ± 0.50 (18) | 22.44 ± 0.58 (10) |
| Tarsus length | 22.88 ± 0.56 (5) | 24.12 ± 0.27 (18) | 23.93 ± 0.31 (10) |
| Scaled mass index | 21.33 ± 1.23 (5) | 21.86 ± 0.47 (18) | 22.39 ± 0.65 (10) |
